# Lrp1 facilitates infection of neurons by Jamestown Canyon virus

**DOI:** 10.1101/2024.11.06.622176

**Authors:** Zachary D. Frey, David A. Price, Kaleigh A. Connors, Rachael E. Rush, Griffin Brown, Cade E. Sterling, Farheen Fatma, Madeline M. Schwarz, Safder Ganaie, Xiaoxia Cui, Zachary P. Wills, Daisy W. Leung, Gaya K. Amarasinghe, Amy L. Hartman

## Abstract

Jamestown Canyon virus (JCV) is a bunyavirus and arbovirus responsible for neuroinvasive disease in the United States. Little is known about JCV pathogenesis, and no host factors required for cellular infection have been identified. Recently, we identified low-density lipoprotein receptor related protein 1 (Lrp1) as a host entry factor for two other bunyaviruses Rift Valley fever virus (RVFV) and Oropouche virus (OROV). Here, we assessed the role of Lrp1 in mediating JCV cellular infection of neurons. Both neuronal and non-neuronal immortalized cell lines deficient for Lrp1 displayed reduction in infection with JCV, and early stages of infection such as binding and internalization were impacted by lack of Lrp1. In primary rat neurons, Lrp1 was highly expressed, and the neurons were highly permissive for JCV infection. Treatment of primary neurons with recombinant receptor-associated protein (RAP), a high affinity ligand for Lrp1, resulted in reduced infectivity with JCV. In addition, pretreatment of cells with RVFV Gn inhibited JCV infection, suggesting that the two viruses may share overlapping binding sites. These results provide compelling evidence that Lrp1 is an important cellular factor for efficient infection by JCV, and thus multiple bunyaviruses with varying clinical manifestations and tissue tropism are facilitated by the host cell Lrp1. Reliance of multiple bunyaviruses on Lrp1 makes it a promising target for pan-bunyaviral antivirals and therapeutics.

## INTRODUCTION

In North America, *Aedes, Culiseta*, and *Anopheles* mosquitoes transmit orthobunyaviruses (order *Bunyavirales*; family *Peribunyaviridae*) including Jamestown Canyon virus (JCV), La Crosse Encephalitis virus (LACV) and California Encephalitis virus (CEV) (1-3). White-tailed deer are the presumed reservoir host of JCV in the United States and Canada, and the high seroprevalence in both deer (∼80%) and humans (up to 20%) in endemic regions highlights the zoonotic potential of this relatively understudied virus (4-7). JCV disease in humans is often asymptomatic or results in a mild febrile illness. However, infection can progress to neuroinvasive disease with symptoms such as encephalitis and meningitis (8, 9). In 2021, JCV was the third most prevalent arbovirus in the United States, and 75% (24/32) of patients infected with JCV were hospitalized leading to two deaths (10). Despite the potential for zoonotic spread and a high rate of hospitalization in reported human cases, there remains a major gap in understanding the capacity for JCV to infect neurons.

The low-density lipoprotein receptor related protein 1 (Lrp1) is an entry factor for two distantly related viruses in *Bunyavirales*: Rift Valley fever virus (RVFV, *Phenuiviridae*) and the recently re-emerged Oropouche orthobunyavirus (OROV, *Peribunyaviridae*) (11, 12). Further, another bunyavirus, Crimean-Congo hemorrhagic fever virus (CCHFV; *Nairoviridae*), uses the related low-density lipoprotein receptor (LDLR) as an entry factor (13-15). Additional RNA viruses were implicated to rely on Lrp1 for later stages of infection, including the peribunyavirus LACV (16). Due to multiple divergent viruses within the order *Bunyavirales* potentially using members of the LDLR family for cellular entry, we screened JCV for Lrp1 dependence for neuronal infection.

Lrp1 is a large (∼600kDa) transmembrane protein that contains an extracellular alpha chain with four ligand-binding clusters, a region of epidermal growth factor repeats, a transmembrane domain, and a cytoplasmic tail. The ligand binding clusters are composed of LDLR class A (LA) repeats, with the clusters I-IV containing 2, 8, 10, and 11 LA repeats, respectively (17). Most ligands for Lrp1 bind to cluster II (CL_II_) and cluster IV (CL_IV_), including the receptor associated protein (RAP) (18). RAP is a molecular chaperone for Lrp1 and other members of the low-density lipoprotein receptor family, and prevents binding of ligands until the receptor localizes to the cell membrane (19). Domains 1 and 3 of RAP (RAP_D3_) can bind to Lrp1, and RAP_D3_ is sufficient to perform the chaperone duties of the full-length protein (20).

Previous studies have shown that the surface glycoproteins of OROV and RVFV bind to CL_II_ and CL_IV_ of Lrp1, with both viruses demonstrating a higher apparent affinity for CL_IV_. OROV and RVFV likely have overlapping binding sites on Lrp1 as a soluble form of RVFV glycoprotein Gn is able to competitively inhibit OROV infection *in vitro* (11, 12).

Here, we found that JCV binding, internalization, and viral production were reduced in cell lines lacking Lrp1. Further, primary neurons were highly permissive to JCV infection, which were found to express stable levels of Lrp1 during ex vivo culture. Using a high affinity ligand for CL_II_ and CL_IV_ of Lrp1, we demonstrate that blocking these regions with murine RAP_D3_ (mRAP_D3_) results in reduced infection in primary neuron cultures. Additionally, pre-treatment with soluble RVFV Gn resulted in a similar reduction in JCV infection, demonstrating that the two viruses likely bind overlapping regions on Lrp1. These findings highlight the role that Lrp1 plays in JCV infection and further underscore Lrp1 as a multi-bunyavirus host factor.

## METHODS

### Cell Lines

Vero E6 (ATCC, CRL-1586) and BV2 (provided by Gaya Amarasinghe) cells were cultured in Dulbecco’s modified Eagle’s medium (DMEM) (ATCC, 30-2002) and supplemented with 1% penicillin/streptomycin (Pen/Strep), 1% L-glutamine (L-Glut), and 10% fetal bovine serum (FBS). N2a (provided by Gaya Amarasinghe) cells were maintained in Eagle’s Minimum Essential Media (EMEM) (ATCC 30-2003) supplemented with 1% Pen/Strep, 1% L-Glut, and 10% FBS. BV2 and N2a Lrp1 KO cell lines were generated and validated as previously described (11) and maintained in the same culture media as their wild type counterparts.

### Virus

The following reagent was obtained through the NIH Biodefense and Emerging Infections Research Resources Repository, NIAID, NIH: Jamestown Canyon virus, 61V-2235, NR-536. Virus was propagated in Vero E6 cells with standard culture conditions using standard D2 media comprised of DMEM supplemented with 1% Pen/Strep, 1% L-Glut, and 2% FBS. A standard viral plaque assay (VPA) was used to determine the infectious titer of the stocks. The agar overlay for the VPA was comprised of 1X minimal essential medium, 2% FBS, 1% Pen/Strep, 1% HEPES buffer, and 0.8% SeaKem agarose (Fisher, BMA50010); the assay incubated at 37° for 3 days, followed by visualization of plaques with 0.1% crystal violet. Passage 1 (p1) from BEI stock was used for the enclosed studies.

### Lrp1 deficient cell line infections

N2a and BV2 cell lines deficient for Lrp1 were previously described and validated (11). Cells were seeded into 24 well plates at 1.25E5 cells/well. On the day of infection, media was removed from each well and replaced with 100 µl of virus diluted to an MOI of 0.1 in standard D2 media. Virus was incubated for an hour rocking every 15 minutes to ensure the monolayer did not dry out. Following the one-hour adsorption period, the inoculum was removed, and the cells were washed once with 1X PBS. Fresh D2 media was added, and the cells incubated for 24 hours prior to sample collection for measurement of viral RNA (vRNA) or infectious titers.

### Binding and Internalization

Lrp1 KO or WT cells were seeded in 24 well plates at 1E5 cells/well. On the day of infection, media was removed and replaced with 200 µl of 10 µM surfen (21) in PBS. Cells were incubated for 30 minutes at 4°C. Following the incubation, surfen solution was removed and replaced with 200 µl of virus diluted to an MOI of 0.1 in standard D2 media. Plates were returned to 4°C for an hour. The inoculum was removed, and cells were washed 5 times with PBS containing 3% bovine serum albumin (BSA) and 0.02% Tween-20. Binding samples were collected by adding 1 mL of Trizol (Fisher, 15-596-018) directly to the cell monolayer. For internalization assays, wells not collected for binding were incubated for one hour in fresh D2 media at 37°C. Cells were washed once with the same wash buffer containing BSA + Tween-20 and samples were collected by adding 1mL of Trizol directly to the cell monolayer.

### Animal Work

Timed-pregnant Long Evans (Crl:LE) rats were purchased from Charles River Laboratories (Wilmington, MA, USA). Fetuses obtained from embryonic day 18 dams were euthanized to obtain the neurons used in this study. All work with animals adhered to *The Guide for the Care and Use of Laboratory Animals* published by the NIH throughout the duration of the study. The University of Pittsburgh is fully accredited by the Association for Assessment and Accreditation of Laboratory Animal Care (AAALAC). The University of Pittsburgh Institutional Animal Care and Use Committee (IACUC) oversaw this work and approved it under protocol number 22051190.

### Primary Neuron Culture

On the day prior to neuron isolation, acid-washed coverslips were coated with PDL/Laminin (Sigma, P7405-5MG; Invitrogen, 23017-015). Dissociation media (DM) comprised of Hanks’ Balanced Salt Solution (Invitrogen, 14175-103) supplemented with 10mM anhydrous MgCl_2_ (Sigma, M8266), 10mM HEPES (Sigma, H3375), and 1mM kynurenic acid was prepared. DM was brought to a pH of 7.2 and sterile filtered prior to use. On the day of isolation, a trypsin solution containing a few crystals of cysteine (Sigma, C7352), 10mL of DM, 4µl 1N NaOH, and 200 units of Papain (Worthington, LS003126); and a trypsin inhibitor solution containing 25mL DM, 0.25g trypsin inhibitor (Fisher, NC9931428), and 10µl 1N NaOH were prepared and filter sterilized. At embryonic day 18, dams were euthanized, and the brains of the embryos were removed and dissected. The cortices were separated from the hippocampus and placed into DM. Five milliliters of trypsin solution was added and cortices were placed in a 37°C water bath for 4 min, swirling occasionally to mix. The trypsin solution was removed, and cortices were immediately washed with trypsin inhibitor once, and then twice more while swirling in the water bath. Following the washes, the trypsin inhibitor was removed and replaced with 5mL of Neurobasal/B27 media, then triturated to dissociate the neurons. The final volume was brought to 10mL of Neurobasal/B27, and cells were counted and plated at a density of 1.5E5 neurons/well for 24-well plates, or 2.5-3E5 neurons/well for 12-well plates. One hour after isolation, the media was removed and replaced with fresh Neurobasal/B27 media. Primary neuron cultures were maintained in Neurobasal/B27 media, which consists of standard Neurobasal Plus Medium (Thermo-Fisher, A3582901) supplemented with 1% Pen/Strep, 1% L-Glut, and 2% B27 Plus Supplement.

### Quantification of viral RNA

RNA isolation was performed using an Invitrogen PureLink RNA/DNA kit (Fisher, 12-183-025) with a modified protocol as previously described (22). Briefly, supernatant was lysed in Trizol (Invitrogen, 15596026) at a dilution of 1:10. Then, 200 µl of chloroform was added to each sample, mixed, and then centrifuged at 12,000 x g for 15 minutes at 4°C to separate the aqueous and organic phases. The aqueous phase was removed and added to an equivalent volume of 70% ethanol. The PureLink RNA kit protocol was then followed for the remainder of the isolation, and RNA was eluted in 40 µl of RNase-free water. RT-qPCR was performed using the SuperScript III Platinum One-Step RT-qPCR Kit (ThermoFisher, 11745-500), following a previously described protocol (22). Primers targeting the JCV L-segment include 5’-CCTAGATGCTCCGTTGTCTATG-3’ (Jamestown-2364For) and 5’-TGCATTATTGGTGTGTGTTTGT-3’ (Jamestown-2448Rev). The Taqman probe used includes (Jamestown-2387 Probe 5’ 6-FAM/ TCAGTACAGTGGGATTAGAAGCTGGGA /BHQ_1 3’).

### Immunofluorescence

Coverslips were fixed and virus inactivated in 4% paraformaldehyde for 15 minutes prior to storage in 1X PBS at 4°C prior to staining. Cells were permeabilized with 0.1% Triton X-100 diluted in 1X PBS for 10 minutes at room temperature. After permeabilization, coverslips were blocked in 5% normal goat serum (ThermoFisher, 50062Z) for an hour at room temperature.

Coverslips were incubated for two hours at room temperature with primary antibodies. Samples were then incubated for an hour with secondary antibodies conjugated to a fluorophore.

Coverslips were counterstained with Hoechst 33258 (Invitrogen, #H1398, 1:1000) and mounted to slides using Gelvatol (provided by the Center for Biologic Imaging). Fluorescent slides were imaged on either Nikon A-1 Confocal at the Center for Biologic Imaging (CBI), or Leica DMI8 inverted fluorescent microscope at the Center for Vaccine Research. Images were processed using Fiji (v1.53). The following antibodies were used for immunofluorescent staining during this study: mouse anti-β III-tubulin (1:500; R&D Systems, MAB1195), custom rabbit anti-JCV N (1:200; Genscript, Y743THG190-16), rabbit anti-Lrp1 (1:500; Abcam, ab92544), antisera from mice immunized with a sublethal dose of JCV (1:200; generated in house), goat anti-rabbit 488 (1:500; Invitrogen, A11008), goat anti-mouse 488 (1:500; Invitrogen, A11001), goat anti-rabbit 594 (1:500; Invitrogen, A11012), and goat anti-mouse 594 (1:500; Invitrogen, A11005).

### Western Blot

Cells were inactivated in 100μl of radioimmunoprecipitation assay buffer (Thermo Fisher Scientific, 89901) with 1% Halt Protease Inhibitor (Thermo Fisher Scientific, 78429) for 10 min at room temperature. Samples were centrifuged at 13,500 relative centrifugal force for 20 min.

Cellular debris was removed, and a bicinchoninic acid (BCA) assay was completed following the manufacturer’s instructions (Thermo Fisher Scientific, Pierce BCA Protein Assay, 23227). Five micrograms of protein from each sample were loaded into a NuPAGE 4 to 12% Bis-Tris gel (Invitrogen, NP0323BOX) and run for 35 min at 165 V. The protein was transferred to a nitrocellulose membrane (LI-COR, 926-31090) using an iBlot 2 system (Invitrogen, IB21001).

Membranes were blocked for 1 hour rocking at room temperature in Intercept® (PBS) Blocking Buffer (Li-Cor, 927-70001). Following the block, membranes were incubated overnight rocking at 4°C with primary antibodies diluted in Intercept® T20 (PBS) Antibody Diluent (Li-Cor, 927-75001). The following primary antibodies were used in this study: mouse anti-GAPDH (1:1000; Invitrogen, MA1-16757), rabbit anti-Lrp1 (1:500; Cell Signaling, 64099S), custom rabbit anti-JCV N (1:500; Genscript, Y743THG190-16), mouse anti-βIII-tubulin (1:500; R&D Systems, MAB1195), anti-RVFV Gn Clone 4D4 (1:500; BEI Resources NR-43190) and mouse anti-β-actin (1:500; Santa Cruz Biotechnology, sc-47778). The following day, the membranes were washed by rocking in 10mL of PBS-T three times for 5 min each. Membranes were probed for 1 hour rocking at room temperature with either goat anti-rabbit IRDye 800CW (1:10,000; Li-Cor, 926-32211), goat anti-rabbit IRDye 680RD (1:10,000; Li-Cor, 925-68071), goat anti-mouse IRDye 800CW (1:10,000; Li-Cor, 925-32210), or goat anti-mouse IRDye 680RD (1:10,000; Li-Cor, 926-68070) diluted in Intercept® T20 (PBS) Antibody Diluent (Li-Cor, 927-75001). The membranes were washed by rocking in 10mL of PBS-T three times for 5 min each, then rinsed with 1X PBS. The membrane was visualized using an Odyssey Clx Imager (LiCor, Lincoln, Nebraska USA).

### Viral Plaque Assay

Vero E6 cells were plated into 12-well plates and allowed to incubate overnight until near confluency. Samples were serially diluted in D2 media. The inoculum was allowed to incubate for one hour at 37°C and then removed. Agar overlay composed of 1X minimal essential medium, 2% FBS, 1% Pen/Strep, 1% HEPES buffer, and 0.8% SeaKem agarose (CAT#s) was added to each well. The assay incubated at 37°C for 3 days to allow for the formation of plaques, fixed with 37% formaldehyde for at least 3 hours, then stained with 0.1% crystal violet for visualization and counting of plaques.

### Viral Growth Curve Infection

Primary rat neurons were maintained in culture for 3 days following isolation. Infection occurred on day 4 in vitro (DIV). JCV was thawed and diluted in D2 media to an MOI of 0.1, 0.01, and 0.001. Media was removed from wells, and 100µl of inoculum was added to each well. Cells were incubated at 37°C for an hour, rocking every 15 minutes to prevent the monolayer from drying out. Following the adsorption period, the inoculum was removed from the wells and replaced with Neurobasal/B27 media. Cells were incubated for 15 minutes, and 100µl of supernatant was inactivated in 900µl of Trizol Reagent (Invitrogen, 15596026) to measure 0hpi viral RNA levels. Timepoint collection of samples occurred at 24, 36, 48, and 60hpi, at which 100µl of supernatant was inactivated in 900µl of Trizol, remaining supernatant was collected and stored at -80°C, and plates were fixed with 4% PFA for 15 minutes and stored at 4°C in 1x PBS for immunofluorescent staining.

### Recombinant Protein Expression and Purification

mRAP_D3_ or mRAP_D3_ (K256A/K270E) expression plasmids were transformed in BL21(DE3) *E. coli* cells (Novagen). Colonies were cultured in Luria Broth media at 37°C to an OD_600_ of 0.6 and induced with 0.5 mM isopropyl-β-D-thiogalactoside (IPTG) for 14 hrs at 18°C. Cells were harvested and resuspended in lysis buffer containing 25 mM sodium phosphate (pH 7.5), 500 mM NaCl, 20 mM imidazole, 5 mM 2-mercaptoethanol, and were lysed using an EmulsiFlex-C5 homogenizer (Avestin). Lysates were clarified by centrifugation at 24,000 x *g* at 4°C for 40 min. Proteins were purified using a series of chromatographic columns as described previously (11). Protein purity was determined by Coomassie staining of SDS-PAGE. Soluble RVFV Gn was expressed in the same manner as mRAP_D3_ and resuspended in a lysis buffer containing 20 mM Tris-HCl (pH 8.0), 500 mM NaCl, 5 mM 2-mercaptoethanol. Following lysis, the Gn pellet was resuspended in 20 mM Tris-HCl (pH 8.0), 500 mM NaCl, 5 mM imidazole, 8 M urea, and 1 mM 2-mercaptoethanol. RVFV Gn was refolded on a NiFF (GE Healthcare) column using a reverse linear urea gradient and eluted with imidazole. Gn_316_ was further purified using a size exclusion column (SD200 10/300L, GE Healthcare).

### Competitive inhibition assays with mRAP_D3_ or RVFV Gn

Primary rat neurons were isolated as described above and maintained in culture for 3 days. Treatment and infection occurred on day 4 in vitro. Proteins were diluted to the desired concentration in both D2 and Neurobasal media. Culture media was partially removed and replaced with Neurobasal containing mRAP_D3_ or Gn. Plates were allowed to incubate for 45 mins at 37°C. Following pre-treatment, all culture media was removed and replaced with D2 containing viral inoculum and either mRAP_D3_ or Gn. Plates were incubated for an hour with rocking every 15 minutes. The inoculum was removed following the adsorption period and Neurobasal containing mRAP_D3_ or Gn was added to the wells and cells returned to the incubator. Twenty-four hours later, supernatant was collected, and plates were fixed with 4% PFA for 15 minutes or cells were lysed with RIPA buffer for 10 minutes. Viral titers were then analyzed by RT-qPCR or VPA and viral antigen was visualized through immunofluorescent staining or Western blot.

### Statistics and Data Analysis

Statistical analysis was performed using GraphPad Prism Version 8.0. Significance was determined by one-way or two-way ANOVA analysis. Error bars show mean and standard deviation. Significance indicated by: *, P<0.05; **, P<0.01; ***, P<0.001; ****, P<0.0001; ns, no significance.

## RESULTS

### Reduced binding and internalization of JCV on cells lacking Lrp1

We previously generated and validated Lrp1 knockout (KO) in both murine N2a (neuroblastoma) and BV2 (microglia) cell lines (11, 12). Using these Lrp1-deficient cells and their Lrp1 sufficient counterparts, we infected each cell type with JCV (strain 61V-2235; MOI=0.1) and measured the amount of viral RNA in the supernatant at 24 hours post-infection (hpi) (**Fig. 1A**). For both N2a and BV2 cells, there was a significant reduction in viral RNA production in the absence of Lrp1. The reduction in infection was visible by immunofluorescence microscopy, where both N2a and BV2 Lrp1 KO cells display decreased staining for JCV nucleoprotein (N) at 24 hpi compared to the wildtype (WT) cells (**Figs. 1C and 1D**).

**Figure 1.**
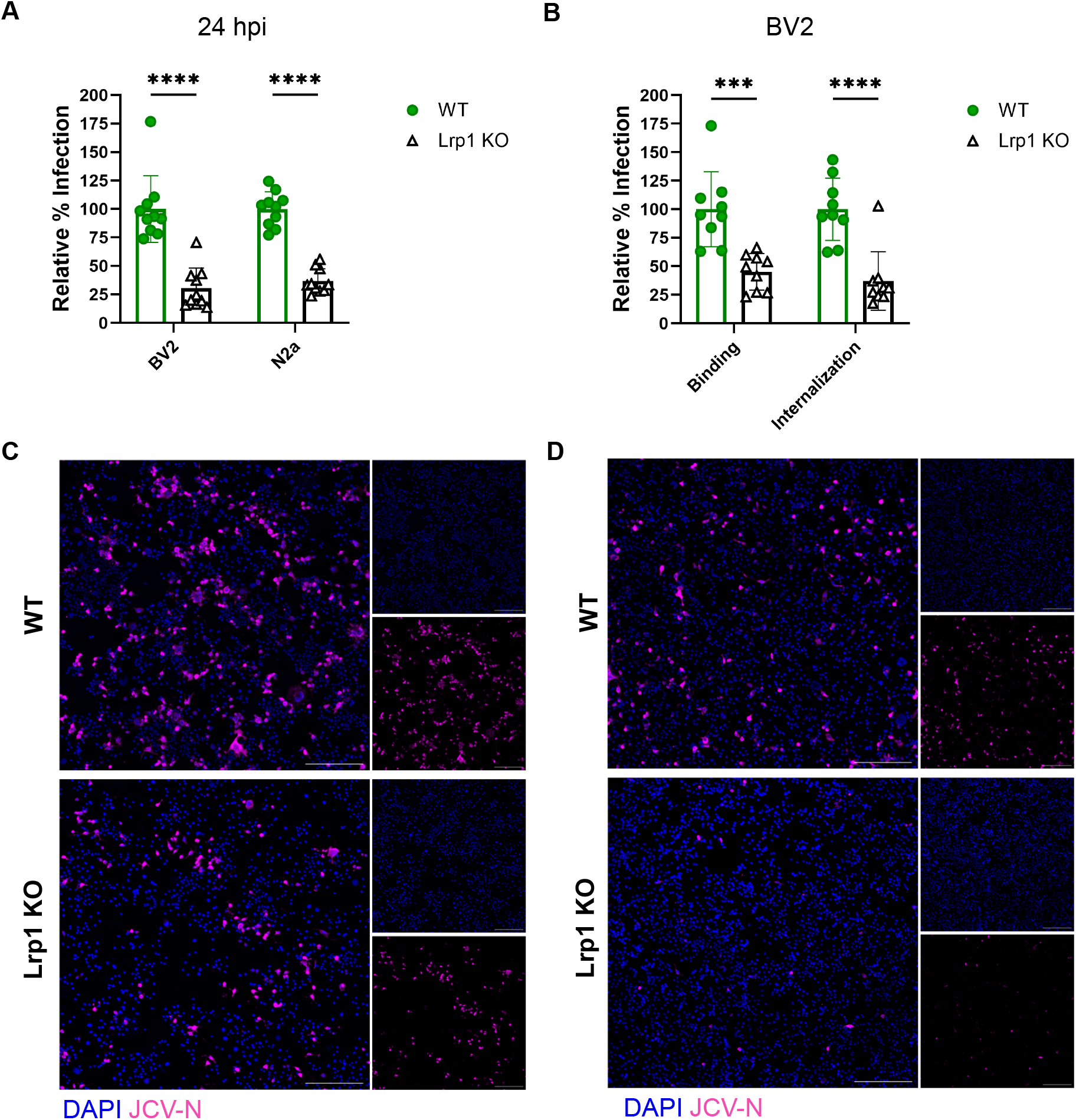
Lrp1 promotes JCV entry. (A) Virus production was determined by infecting cells at an MOI of 0.1 and quantifying viral RNA from the supernatant at 24hpi. (B) Wild type and Lrp1 knockout BV2 cells were incubated with surfen at 4°C for 30 minutes. Surfen was removed and cells were incubated with virus (MOI 0.1) for 1 hour at 4°C. Cells were washed and binding samples were collected. Cells were returned to 37°C for 1 hour and internalization samples were collected. At 24hpi, N2a (C) and BV2 (D) cells were fixed with 4% PFA and stained for JCV-N (pink) and counterstained with Hoescht (blue). Slides were imaged at 10X using a Leica DMI8 inverted microscope. Scale bar = 250µm. Statistics determined by two-way ANOVA. *** p=0.0002, **** p<0.0001.

To determine the effect of Lrp1 on JCV binding and internalization, we used BV2 cells. Cells were first treated with the glycosaminoglycan (GAG) antagonist surfen to prevent any non-specific binding to proteoglycans (21), incubated with JCV (MOI=0.1) for 1 hr at 4°C to allow binding but not internalization, and washed extensively before collection and RNA quantification. For internalization assays, cells were incubated at 37°C for another 1 hour after washing. We observed a 50-60% reduction in both binding and internalization in BV2 Lrp1 KO cells when compared to the WT cells (**Fig. 1B**). These results indicate that Lrp1 is utilized for efficient JCV binding and internalization, and this defect persists through to 24 hpi.

### Primary neurons are permissive to JCV Infection

Given that both N2a and BV2 cells are immortalized cell lines, and that little is known about JCV replication in neurons, we isolated primary rat neurons to study the interaction between Lrp1 and JCV. Primary cortical neurons from rat embryos were isolated and cultured for 4 days in vitro (DIV). We initially conducted growth curves by infecting neurons with JCV at MOIs of 0.1, 0.01, and 0.001 and assessing viral production over time. Supernatants were analyzed for viral RNA (RT-qPCR) and viral plaque assay (VPA) for infectious titers. Infection of primary neuron cultures with JCV resulted in high levels of virus production in a dose-dependent manner (**Fig. 2A**). Within 24 hours, viral RNA reached levels between 1E4 to 1E6 plaque forming unit (PFU) equivalents (eq.)/mL. By 60 hpi, all MOIs reached 1E6 PFU/mL or PFU eq./mL (**Fig. 2A, Supplemental Fig. 1A**). Viral infection was visualized in the neurons via immunofluorescence microscopy by staining for JCV N antigen and the neuronal marker βIII-Tubulin (**Fig. 2B, Supplemental Fig. 1B**). Mock-infected cultures were stained with βIII-Tubulin appeared healthy containing neurons with long cellular processes. At an MOI of 0.1, JCV antigen staining was widespread throughout the cultures at 24 hpi and remained prevalent at 48 hpi. As the infection progressed, the cellular debris in culture increased resulting in a punctate βIII-Tubulin staining pattern, indicating loss of neuronal structure (**Fig. 2B, Supplemental Fig. 1B**). Under the culture conditions used here, neurons expressed Lrp1 throughout the culture period (4 to 7 DIV) (**Fig. 2C**). Lrp1 expression was widely detectable by microscopy and was found in both the processes and cell bodies (**Fig. 2D**; **Supplemental Fig. 1C**).

**Figure 2.**
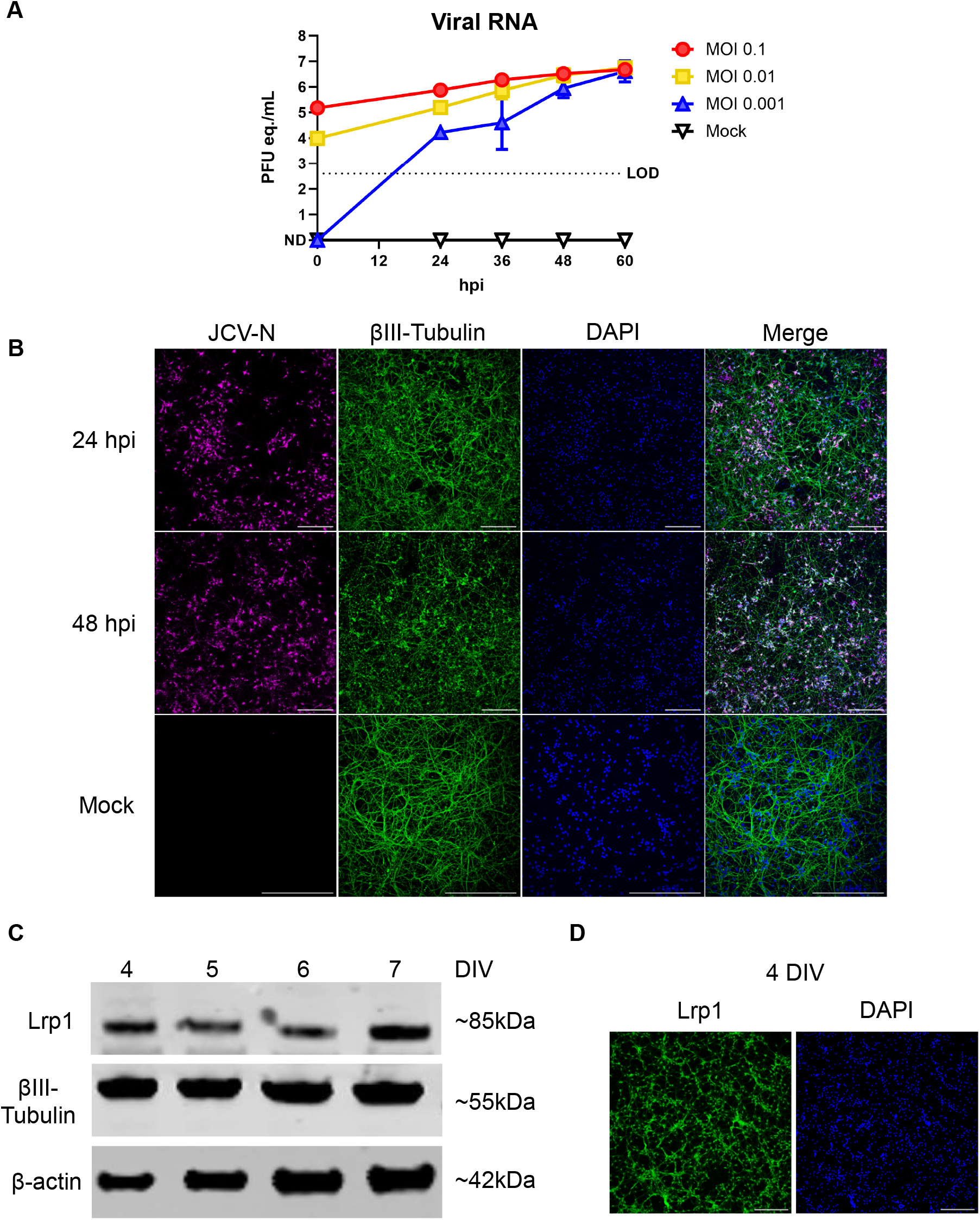
Replication kinetics of JCV in primary rat neurons. Primary rat neurons were infected with JCV at an MOI of 0.1, 0.01, or 0.001. (A) Viral RNA was quantified at 24, 36, 48, and 60 hpi timepoints. (B) Infected or mock infected coverslips were fixed in 4% PFA and stained for JCV-N (pink) and βIII-Tubulin (green) and counterstained with Hoescht (blue). Slides were imaged at 20X using a Nikon A-1 confocal microscope. Scale bar = 250µm. (C) Western blot of uninfected primary rat neurons across different days in vitro (DIV). Blots were probed for the 85 kDa beta chain of Lrp1, βIII-Tubulin, and β-Actin. (D) Immunofluorescent microscopy of neurons 4 DIV. Coverslips were fixed with 4% PFA and stained for Lrp1 (green) and counterstained with Hoescht (blue). Slides were imaged at 10X using a Leica DMI8 inverted microscope. Scale bar = 250µm.

### Treatment of primary neuron cultures with a high-affinity Lrp1 binding protein reduces JCV Infection

Receptor associated protein (RAP) is an intracellular high affinity Lrp1 chaperone protein known to competitively inhibit ligand binding to the CL_II_ and CL_IV_ domains of Lrp1 (18). Domain 3 of the mouse RAP protein (mRAP_D3_) can be added exogenously to cells prior to infection to interrogate the dependence on Lrp1 for infection, as we demonstrated with RVFV and OROV (11, 12). Here, primary neurons were pre-treated with recombinant mRAP_D3_ or a mutated version of mRAP_D3_ containing K256A/K270E mutations which causes a reduced affinity for Lrp1 (11, 23), followed by infection with JCV (MOI 0.1). At 24 hpi, viral RNA levels in the supernatant were reduced approximately 75-90% in a dose-dependent manner compared to the infected untreated controls (**Fig. 3A**). The mutant mRAP_D3_, in comparison, was not as effective at reducing JCV viral RNA, and reached a similar decrease in RNA titers (∼75%) only at the highest dose tested (10µg/mL) (**Supplemental Fig. 2A**). Plaque assays measuring infectious titers at 24 hpi showed a similar reduction to viral RNA after mRAP_D3_ treatment (**Fig. 3B**). By microscopy, viral antigen in mRAP_D3_-treated cells was restricted to small foci as opposed to being widespread throughout the culture in the untreated control images (**Fig. 3C, Supplemental Fig. 2B**). Thus, exogenous mRAP_D3_ can inhibit JCV infection in primary rat neurons through competition for binding to Lrp1.

**Figure 3.**
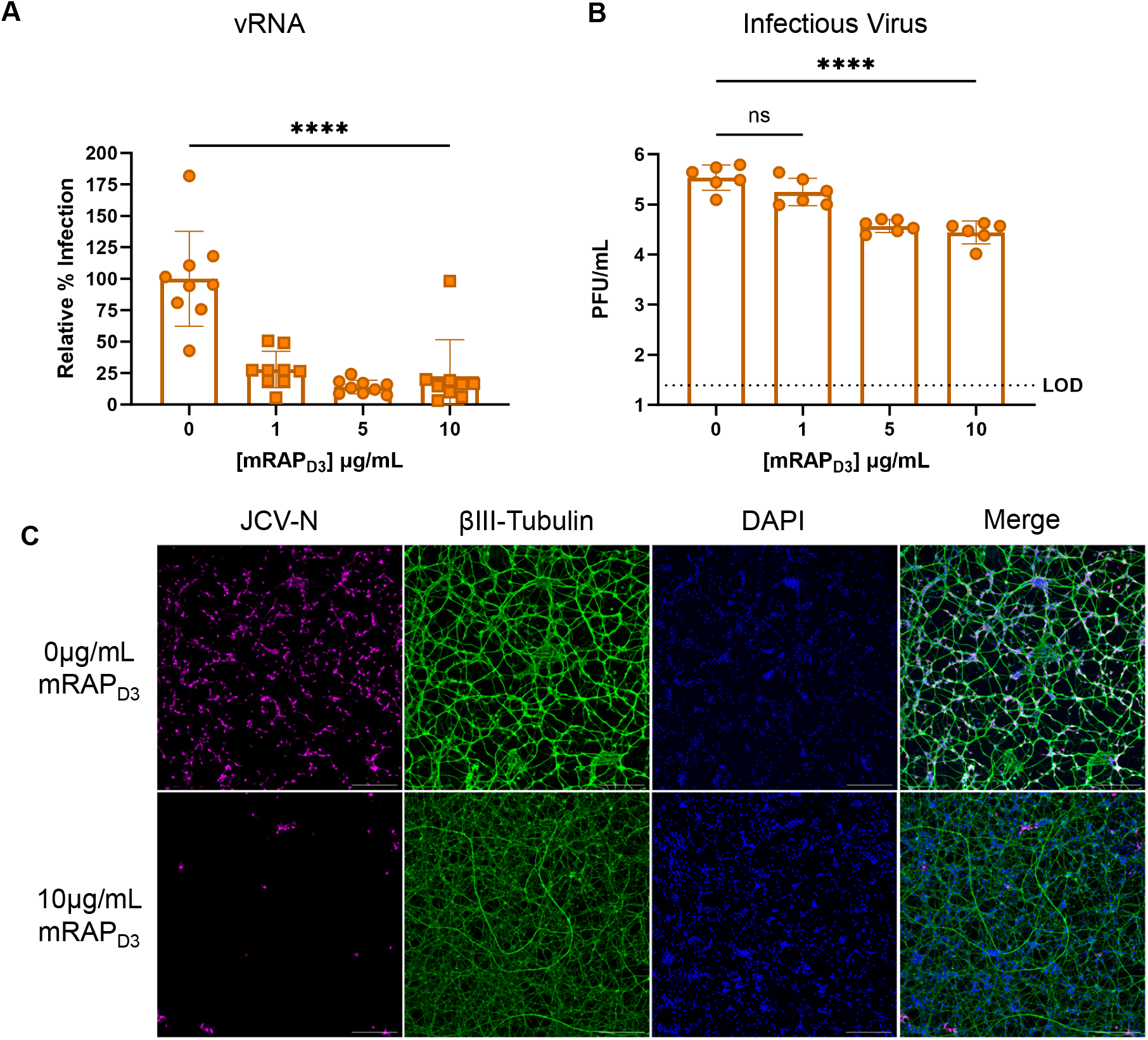
Pre-treatment with a high affinity Lrp1 binding protein reduces JCV infection. Primary rat neurons were pre-treated with different concentrations of mRAP_D3_ for 45 minutes followed by infection with JCV at an MOI of 0.1. At 24hpi, supernatant was collected for quantification of (A) viral RNA and (B) infectious virus. (C) Coverslips were fixed with 4% PFA and stained for JCV-N (pink) and βIII-Tubulin (green) and counterstained with Hoescht (blue). Slides were imaged at 10X using a Leica DMI8 inverted microscope. Scale bar = 250µm. Statistics determined by one-way ANOVA. **** p<0.0001.

### Exogenous Gn protein from RVFV restricts JCV infection of primary neurons

The Gn glycoprotein of the distantly related bunyavirus RVFV binds to CL_II_ and CL_IV_ of Lrp1, and exogenous treatment of cells with recombinant RVFV Gn competitively inhibited both homologous infection with RVFV and heterologous infection by OROV (11, 12). In a similar heterologous competition experiment to further probe the role of Lrp1 in JCV infection, primary neurons were pre-treated with recombinant RVFV Gn for an hour, followed by infection with JCV at an MOI 0.1. At 24 hpi, viral titer was measured by RT-qPCR. JCV titers were significantly reduced in the presence of 5 and 10 µg/mL of exogenous RVFV Gn (**Fig. 4A**). By western blot, as RVFV Gn levels increased, the amount of JCV N protein detected in culture lysates decreased (**Fig. 4B**). Immunofluorescence microscopy revealed a decrease in viral antigen staining in cells treated with RVFV Gn compared to untreated cells (**Fig. 4C, Supplemental Fig. 3A**). Our results showing that RVFV Gn can competitively inhibit and reduce JCV infection suggests that JCV likely binds an overlapping binding site on Lrp1 CL_II_ and CL_IV_.

**Figure 4.**
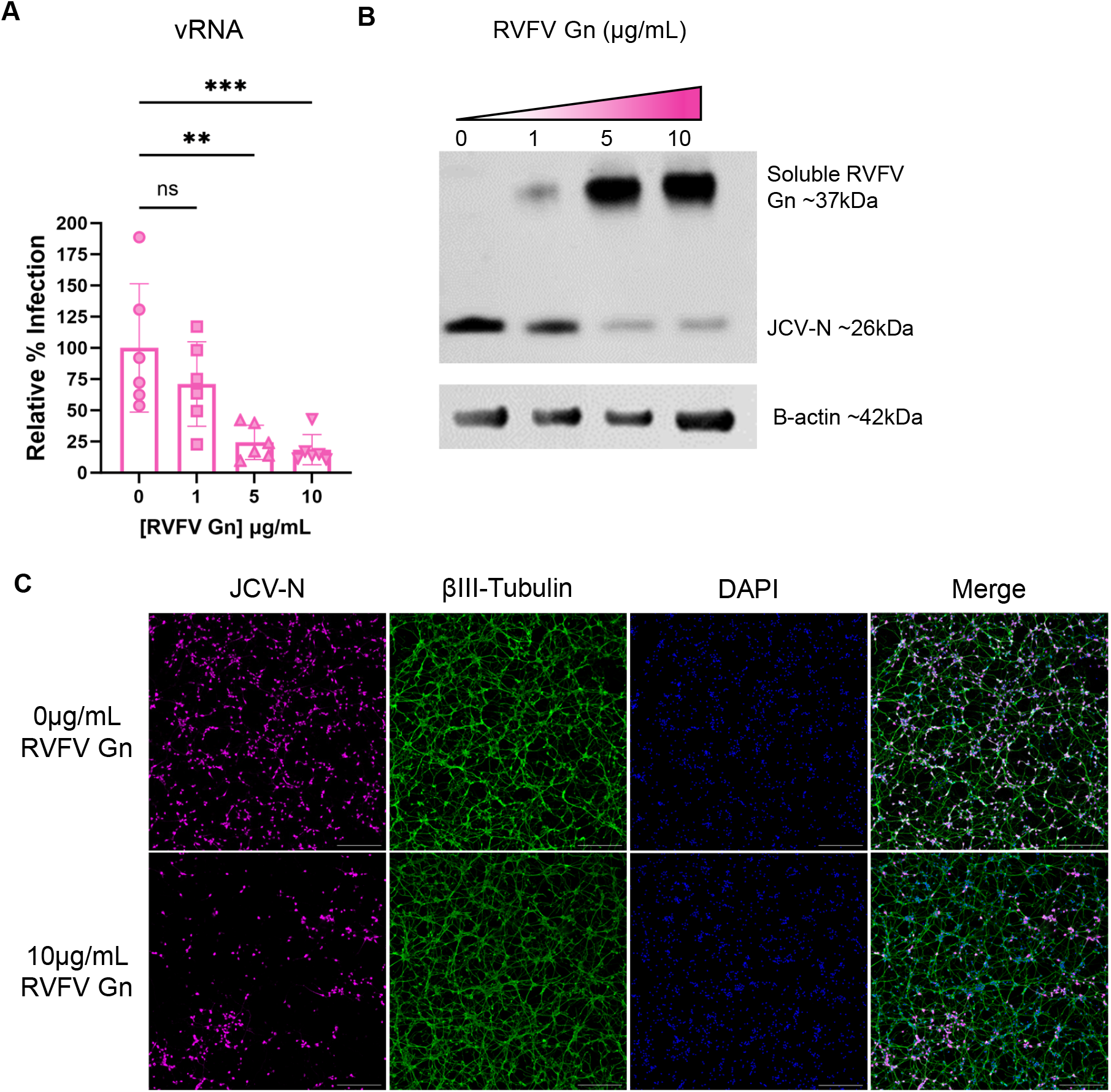
Pre-treatment with RVFV Gn reduces JCV infection. Primary rat neurons were pre-treated with different concentrations of recombinant, soluble RVFV Gn for 45 minutes followed by infection with JCV at an MOI of 0.1. At 24hpi, supernatant was collected for quantification of (A) viral RNA. Cells were lysed in RIPA buffer and to assess protein levels via Western blot (B). Western blots were probed for RVFV Gn, JCV-N, and β-actin. (C) Coverslips were fixed with 4% PFA and stained for JCV-N (pink) and βIII-Tubulin (green) and counterstained with DAPI (blue). Slides were imaged at 10X using a Leica DMI8 inverted microscope. Scale bar = 250µm. Statistics determined by one-way ANOVA. **p=0.0016, ***p=0.0008.

## DISCUSSION

Jamestown Canyon virus is a prevalent arbovirus found in white-tailed deer populations in North America. While severe disease in people may be rare compared to overall seropositivity rates, the potential for further spread given deer-human proximity and the capacity to induce severe neurological clinical outcomes makes JCV an arbovirus of concern for the U.S. and Canada (4-7). Surprisingly, little experimental work has been conducted to determine its tropism for the central nervous system. Immunocompetent mice have previously been used to study JCV neuropathogenesis; however, lack of neuroinvasion make studying virus-cell interactions in the brain challenging (24, 25). Intranasal and intracranial inoculation of JCV results in consistent neurologic disease in mice (24, 26), but this does not mimic a natural infection route as JCV is primarily spread by mosquitos. Mice deficient in type I interferon receptors or key signaling molecules (IRF3, IRF7, or MAVS) develop neurologic disease following intraperitoneal infection, demonstrating that innate immunity is responsible for controlling JCV in the periphery and preventing neuroinvasion (27).

Here, we used primary rat neurons as a relevant ex vivo primary cell model to study JCV neuropathogenesis, as rat neurons can be obtained relatively easily in large numbers, and our lab has experience using a rat model to study viral encephalitis (28-33). After several days of culture, the isolated cells displayed phenotypic characteristics of neurons including long processes and intense staining with the neuronal marker βIII-tubulin. The neurons were highly permissive to JCV infection, with robust replication of JCV within 24 hours after infection.

Extensive visual cytopathic effect was evident by loss of neuronal processes and accumulation of cellular debris within the cultures. A previous study that used the neuronal cell line SH-SY5Y and human neural stem cells (hNSCs) to study JCV replication in vitro found that JCV replicates slower and causes less cytopathic effect when compared to other California Serogroup viruses (24). As primary rat neurons showed robust replication and extensive cytopathic effect quickly after infection, they may serve as an attractive alternative to cell lines for studying JCV in vitro.

The low-density lipoprotein receptor (LDLR) family of cell surface receptors is an evolutionarily conserved family of proteins with a variety of functions, including lipoprotein metabolism and cellular signaling (34). These molecules have been implicated in mediating cellular entry of a variety of arboviruses, including multiple alphaviruses and bunyaviruses (11-16, 35-38). Many of these viruses have a wide host range and tissue tropism, which is supported by the evolutionary conservation and broad tissue distribution of the LDLR family members. Lrp1 differs from other LDLRs that serve as viral receptors, such as LDLR, VLDLR, and ApoER2, in that it contains four ligand binding cluster domains, while the other smaller family members are comprised of just one (39). This enables Lrp1 to interact with ligands through multiple clusters, differentiating its interactions with ligands from the smaller members of the LDLR family (40). RVFV and OROV infection are supported by binding to CL_II_ and CL_IV_ of Lrp1 (11, 12), and it is possible that both clusters interact with the virion during the course of infection, complicating the molecular interactions between viruses and Lrp1.

Neurons and other cells of the CNS express Lrp1 (41) and Lrp1 has a variety of critical functions in the brain including the modulation of NMDA receptor signaling (42), neuronal glucose metabolism (43), and AMPA receptor stability (44). Lrp1 has also been implicated in multiple neurodegenerative diseases, including Alzheimer’s disease, Parkinson’s disease, and Lewy body dementia (45-48). Other LDLRs also play important and often overlapping roles in the CNS. VLDLR and ApoER2 have been found to modulate synaptic plasticity (49) and neuronal migration (50). Mice with Lrp1 deleted on a majority of their neurons (Lrp1^f/f^ Synapsin-Cre) display deficits in motor function (51), and VLDLR and ApoER2 double knockout mice display progressive hind limb paralysis and smaller brain size when compared to WT mice (50), demonstrating the importance of LDLR family members in the CNS function.

Given the conserved use of Lrp1 by distantly related bunyaviruses RVFV and OROV, and given the importance of Lrp1 expression in neurons, we interrogated the dependence on Lrp1 for infection of neurons by JCV. Initial studies screened multiple murine cell lines clonally KO for Lrp1, and we found reduced JCV binding and internalization in the absence of Lrp1. This implicates Lrp1 in the entry stage of infection, similar to its apparent role in infection with RVFV and OROV. We further probed the reliance on Lrp1 using an ex vivo primary rat neuron model in combination with the previously described molecular and biochemical tools of mRAP_D3_ and recombinant RVFV Gn (11, 12). Pre-treatment of primary neurons with either mRAP_D3_ or recombinant Gn from RVFV reduced infection of primary rat neurons. As both mRAP_D3_ and RVFV Gn bind to CL_II_ and CL_IV_ (11, 18), these regions are likely involved with JCV infection.

Additionally, the fact that RVFV Gn can inhibit JCV infection implies that both viruses may use overlapping regions on Lrp1.

Limitations to this study include the fact that we are not able to completely prevent JCV infection by blocking or knocking out Lrp1, suggesting that there are other attachment factors and/or receptors that JCV is able to use for entry. Further, there may be mechanisms of non-specific viral uptake. One such possibility is the use of heparan sulfate, which facilitates entry of the related bunyaviruses RVFV, LACV, Schmallenberg virus, and Akabane virus (52-54). It is possible then that JCV may also use this proteoglycan for attachment and entry. Another possibility is DC-SIGN, which has been implicated as a receptor for RVFV and LACV (55, 56). While DC-SIGN is not known to be expressed by neurons, it is expressed by microglia (57) and dendritic cells (58), and therefore could have an impact on neuropathogenesis. It is also possible that JCV uses an alternative receptor or attachment factor yet to be identified.

While the work presented here strongly suggests that Lrp1 supports efficient infection by JCV, future studies will investigate the direct mechanism of the interaction between Lrp1 and the surface glycoproteins Gn/Gc of JCV. This is a necessary next step to definitively demonstrate that Lrp1 is mediating entry through direct interaction with the JCV surface glycoproteins. While RVFV binds to Lrp1 through interactions with the surface glycoprotein Gn (11), there are large differences in the glycoprotein structures of viruses within *Bunyavirales* (59). Additionally, Crimean-Congo hemorrhagic fever virus, a more distantly related bunyavirus in *Nairoviridae*, interacts with its receptor, low density lipoprotein receptor (LDLR), through the Gc glycoprotein (14, 15). Therefore, it is likely that JCV may engage Lrp1 in a different manner than RVFV does, including potential binding by Gc rather than Gn.

In summary, we present evidence that Lrp1 is a host factor involved in the early stages of neuronal entry by JCV. Based on our previous work with RVFV and OROV, and this study with JCV, Lrp1 is implicated as a multi-bunyaviral host factor. The fact that Lrp1 is highly conserved and is utilized in early infection of diverse bunyaviruses make it an attractive target for the development of broad bunyavirus therapeutics.

## ACKNOWLEDGEMENTS

The following reagent was obtained through BEI Resources, NIAID, NIH: Jamestown Canyon Virus, 61V-2235, NR-536. The following reagent was obtained from the Joel M. Dalrymple – Clarence J. Peters USAMRIID Antibody Collection through BEI Resources, NIAID, NIH: Monoclonal Anti-Rift Valley Fever Virus Gn Glycoprotein, Clone 4D4 (produced *in vitro*), NR-43190.

## FIGURE LEGENDS

**Supplemental Figure 1.**
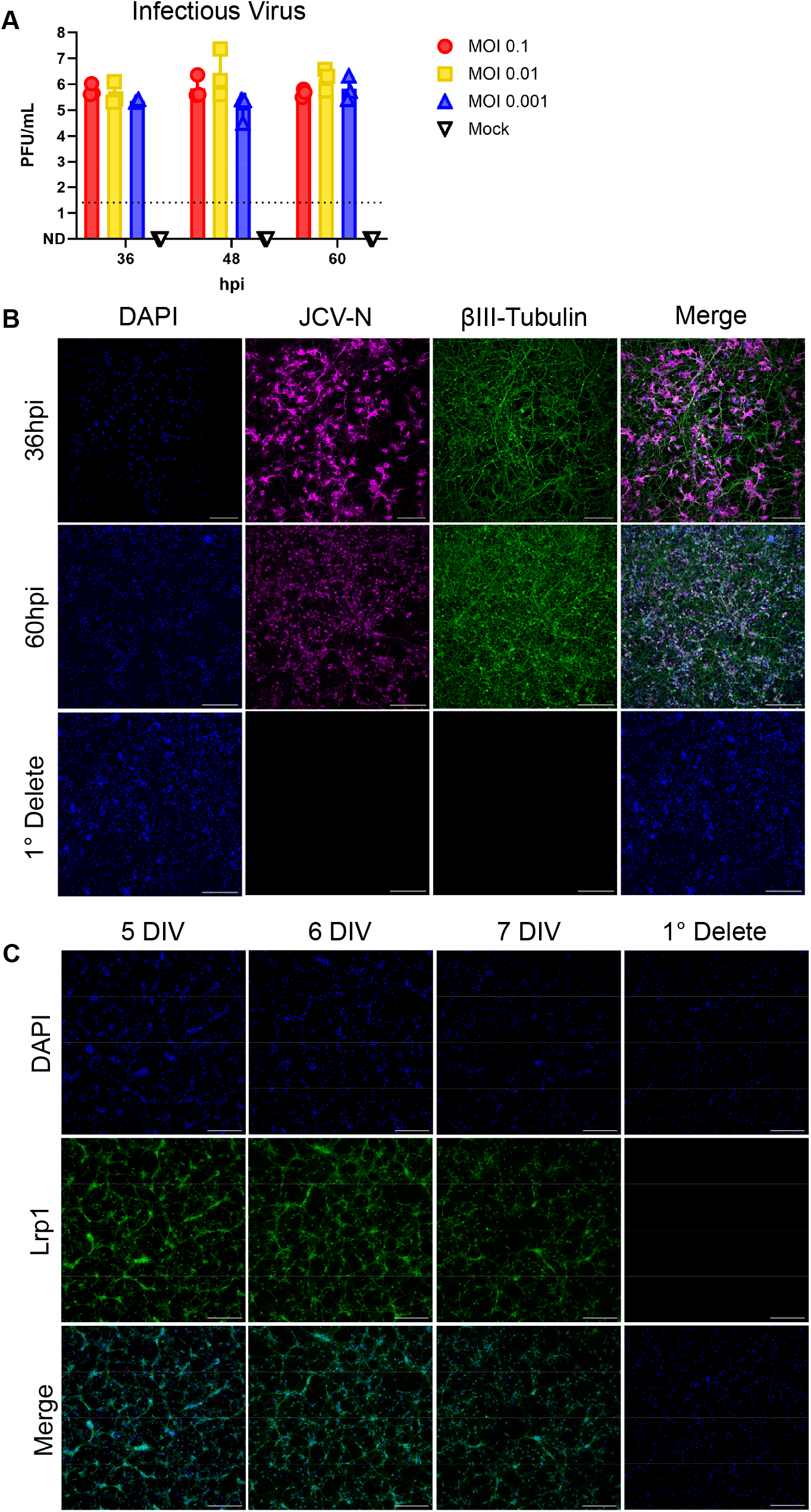
Infectious titers and additional images of immunofluorescent microscopy and from Fig. 2. (A) Infectious titers at 36, 48, 60 hpi. (B) Additional images of 36 hpi, 60 hpi, and primary delete of JCV infected primary rat neurons. Coverslips were stained for JCV-N (pink) and βIII-Tubulin (green) and counterstained with DAPI (blue). Slides were imaged at 20X using a Nikon A-1 confocal microscope. Scale bar = 250µm. (C) Additional images showing Lrp1 expression in primary rat neurons across days 5, 6, and 7 in vitro. Coverslips were stained for Lrp1 (green) and counterstained with DAPI (blue). Slides were imaged at 10X using a Leica DMI8 inverted microscope. Scale bar = 250µm.

**Supplemental Figure 2.**
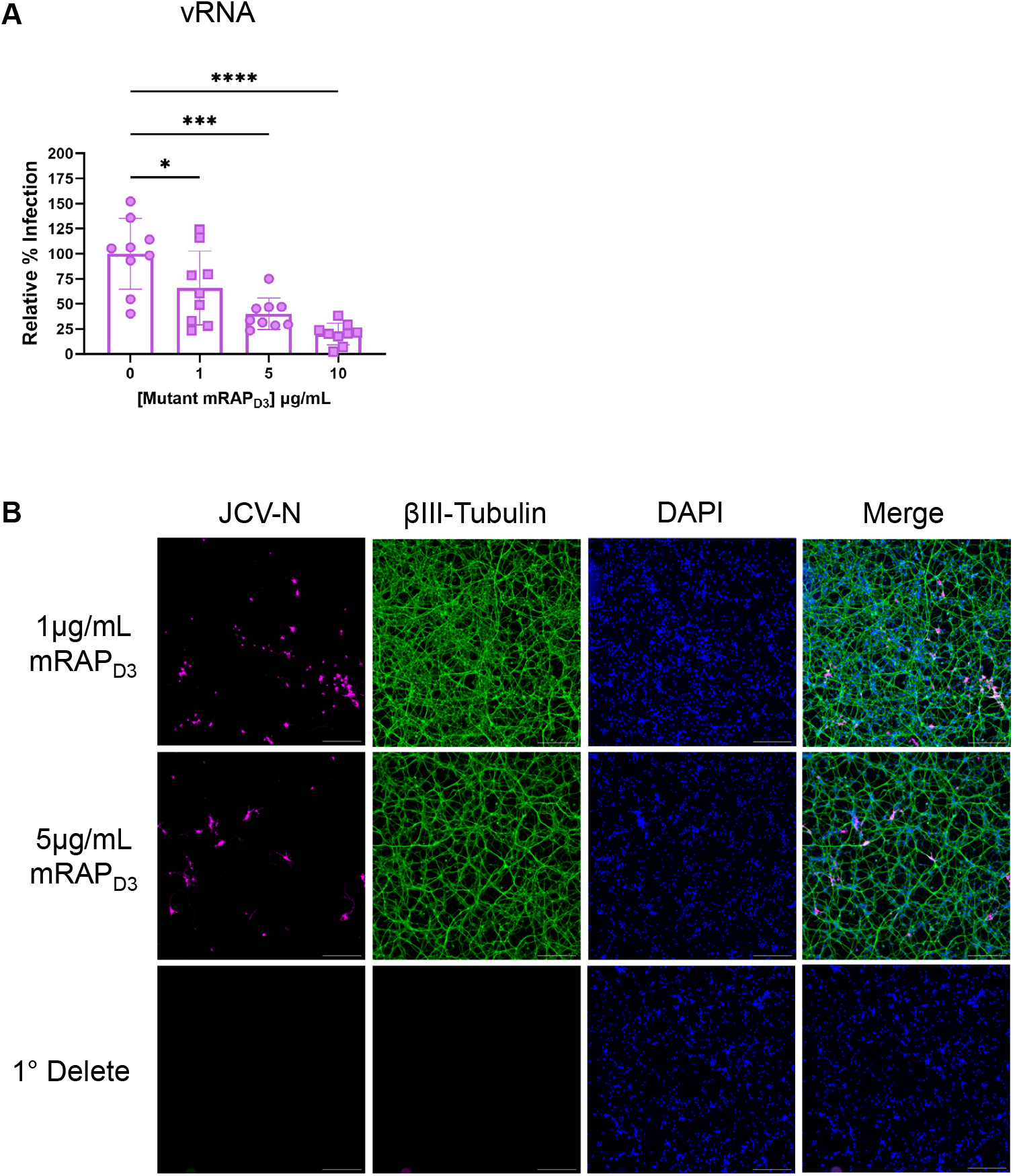
Mutant mRAP_D3_ treatment and additional images of immunofluorescent microscopy from Figure 3. (A) vRNA from K256A/K270E mutant mRAP_D3_ treated cells. (B) Additional images of mRAP_D3_ treatment of primary rat neurons, including primary delete. Coverslips were stained for JCV-N (pink) and βIII-Tubulin (green) and counterstained with Hoescht (blue). Slides were imaged at 10X using a Leica DMI8 inverted microscope. Scale bar = 250µm. Statistics determined by one-way ANOVA. *p=0.0315, ***p=0.0001, ****p<0.0001.

**Supplemental Figure 3.**
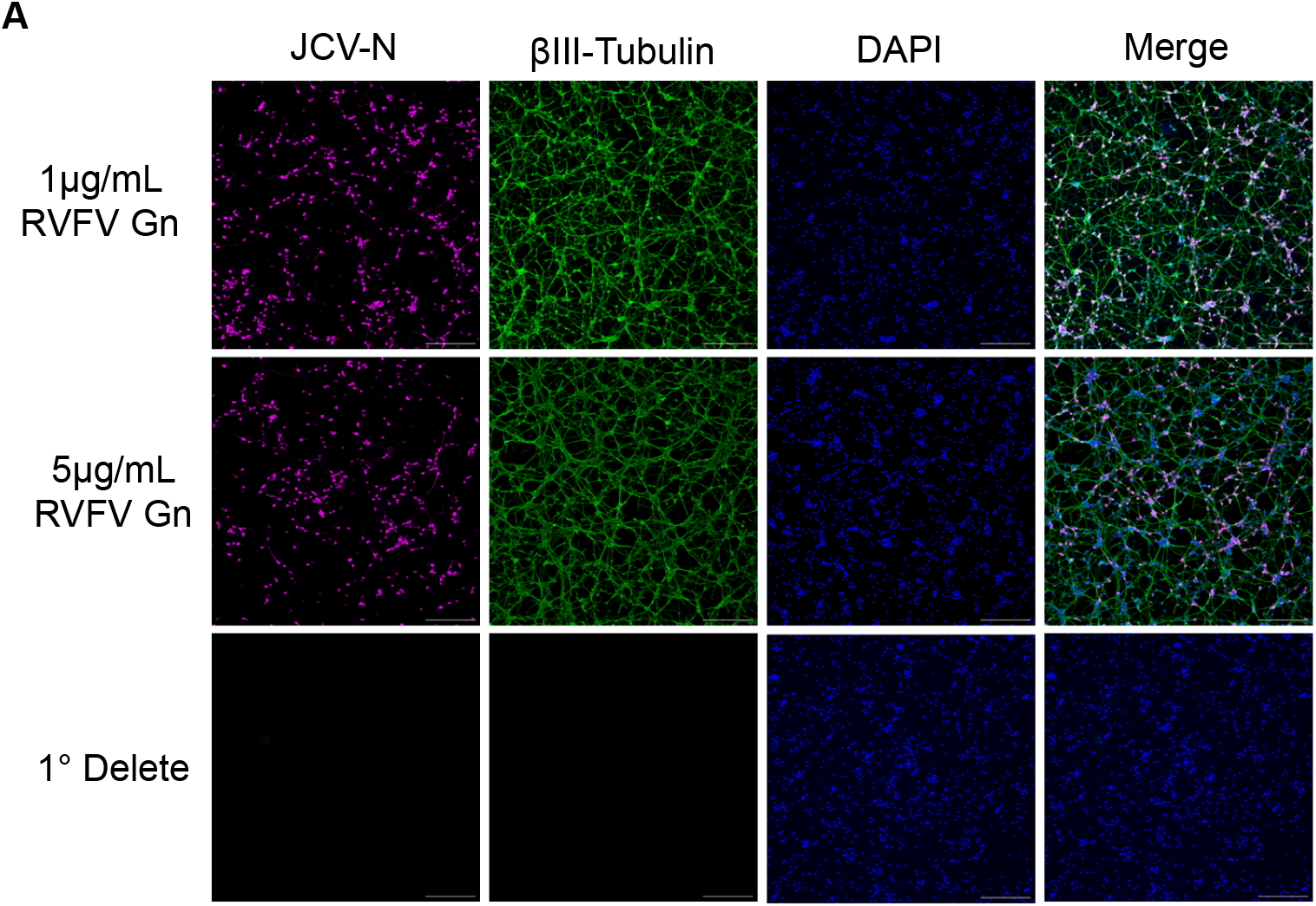
Additional images of immunofluorescent microscopy from Figure 4. (A) Additional images of RVFV Gn treatment of primary rat neurons, including primary delete. Coverslips were stained for JCV-N (pink) and βIII-Tubulin (green) and counterstained with Hoescht (blue). Slides were imaged at 10X using a Leica DMI8 inverted microscope. Scale bar = 250µm.

